# Reconstructing a metazoan genetic pathway with transcriptome-wide epistasis measurements

**DOI:** 10.1101/112920

**Authors:** David Angeles-Albores, Carmie Puckett Robinson, Brian A. Williams, Barbara J. Wold, Paul W. Sternberg

## Abstract

RNA-seq is commonly used to identify genetic modules that respond to perturbations. In single cells, transcriptomes have been used as phenotypes, but this concept has not been applied to whole-organism RNA-seq. Linear models can quantify expression effects of individual mutants and identify epistatic effects in double mutants. To make interpretation of these high-dimensional measurements intuitive, we developed a single coefficient to quantify transcriptome-wide epistasis that accurately reflects the underlying interactions. To demonstrate our approach, we sequenced four single and two double mutants of *Caenorhabditis elegans*. From these mutants, we reconstructed the known hypoxia pathway. In addition, we uncovered a class of 56 genes that have opposing changes in expression in *egl-9(lf)* compared to *vhl-1(lf)* but the *egl-9(lf); vhl-1(lf)* mutant has the same phenotype as *egl-9(lf)*. This class violates the classical model of HIF-1 regulation, but can be explained by postulating a role of hydroxylated HIF-1 in transcriptional control.

**Significance Statement:** Transcriptome profiling is a way to quickly and quantitatively measure gene expression level. Because of their quantitative nature, there is widespread interest in using transcriptomic profiles as a phenotype for genetic analysis. However, a source of major concern is that whole-animal transcriptomic profiles mix the expression signatures of multiple cellular states, making it hard to accurately reconstruct genetic interactions. Additionally, it has been difficult to quantify epistasis, the signature of genetic interaction between two genes, in these molecular phenotypes. Here, we show that it is possible to accurately reconstruct genetic interactions between genes using whole-animal RNA sequencing, and we demonstrate a powerful new way to measure and understand epistasis arising from these measurements. This suggests that whole-organism RNA-seq can be a powerful tool with which to understand genetic interactions in entire organisms and not only in isolated cells. With the advent of genome engineering tools, generating mutants has become easier and faster for many organisms. As mutants become easier to create, phenotyping them has become a major bottleneck in understanding the biological functions of the genes in question. Our work presents a possible solution to this problem, because transcriptome profiling is fast and sensitive to genetic perturbations regardless of the context they operate in.

## Introduction

Genetic analysis of molecular pathways has traditionally been performed through epistatic analysis. Generalized epistasis indicates that two genes interact functionally; such interaction can involve the direct interaction of their products or the interaction of any consequence of their function^1^. If two genes interact, and the mutants of these genes have a quantifiable phenotype, the double mutant of interacting genes will have a phenotype that is not the sum of the phenotypes of the single mutants. Epistasis analysis remains a cornerstone of genetics today^2^.

Recently, biological studies have shifted in focus from studying single genes to studying all genes in parallel. In particular, RNA-seq^3^ enables biologists to identify genes that change expression in response to a perturbation. RNA-seq has been successfully used to identify genetic modules involved in a variety of processes, such as in the *Caenorhabditis elegans* linker cell migration^4^, and planarian stem cell maintenance^5,6^. For the most part, the role of transcriptional profiling has been restricted to target gene identification, and so far there are only a few examples where transcriptomes have been used to generate quantitative genetic models of any kind. In population genetics, eQTL studies have established the power of transcriptomes for genetic mapping^7,8,9,10^. Genetic pathway analysis via epistasis has only been performed once in *Saccharomyces cerevisiae*^11^ and once in *Dictyostelium discoideum*^12^. Recently, Dixit *et al* described a protocol for epistasis analysis in T-cells using single-cell RNA-seq^13^. Epistasis analysis of single cells or single-celled organisms is popular because of the concern that whole-organism sequencing will mix information from multiple cell types, preventing the accurate reconstruction of genetic interactions. Using whole-organism transcriptome profiling, we have recently identified a new developmental state of *C. elegans* caused by loss of a single cell type (sperm cells)^14^, which suggests that whole-organism transcriptome profiling contains sufficient information for epistatic analysis. To investigate the ability of whole-organism transcriptomes to serve as quantitative phenotypes for epistatic analysis in metazoans, we sequenced the transcriptomes of of four well-characterized loss-of-function mutants in the *C. elegans* hypoxia pathway^15,16,17,18^.

Metazoans depend on the presence of oxygen in sufficient concentrations to support aerobic metabolism. Hypoxia inducible factors (HIFs) are an important group of oxygen-responsive genes that are highly conserved in metazoans^19^. A common mechanism for hypoxia-response induction is heterodimerization between a HIF*α* and a HIF*β* subunit; the heterodimer then initiates transcription of target genes^20^. The number and complexity of HIFs varies throughout metazoans. In the roundworm *C. elegans* there is a single HIF*α* gene, *hif-1*^18^, and a single HIF*β* gene, *ahr-1*^21^.

Levels of HIF*α* proteins are tightly regulated. Under conditions of normoxia, HIF-1*α* exists in the cytoplasm and partakes in a futile cycle of protein production and rapid degradation^22^. In *C. elegans*, HIF-1*α* is hydroxylated by a proline hydroxylase (EGL-9)^23^. HIF-1 hydroxylation increases its binding affinity to Von Hippel-Lindau tumor suppressor 1 (VHL-1), which in turn allows ubiquitination of HIF-1 leading to its subsequent degradation. In *C. elegans*, EGL-9 activity is inhibited by binding of CYSL-1, a homolog of sulfhydrylases/cysteine synthases; and CYSL-1 activity is in turn inhibited by the putative transmembrane O-acyltransferase RHY-1, possibly by post-translational modifications to CYSL-1^24^ (see Fig. 1).

**Figure 1.**
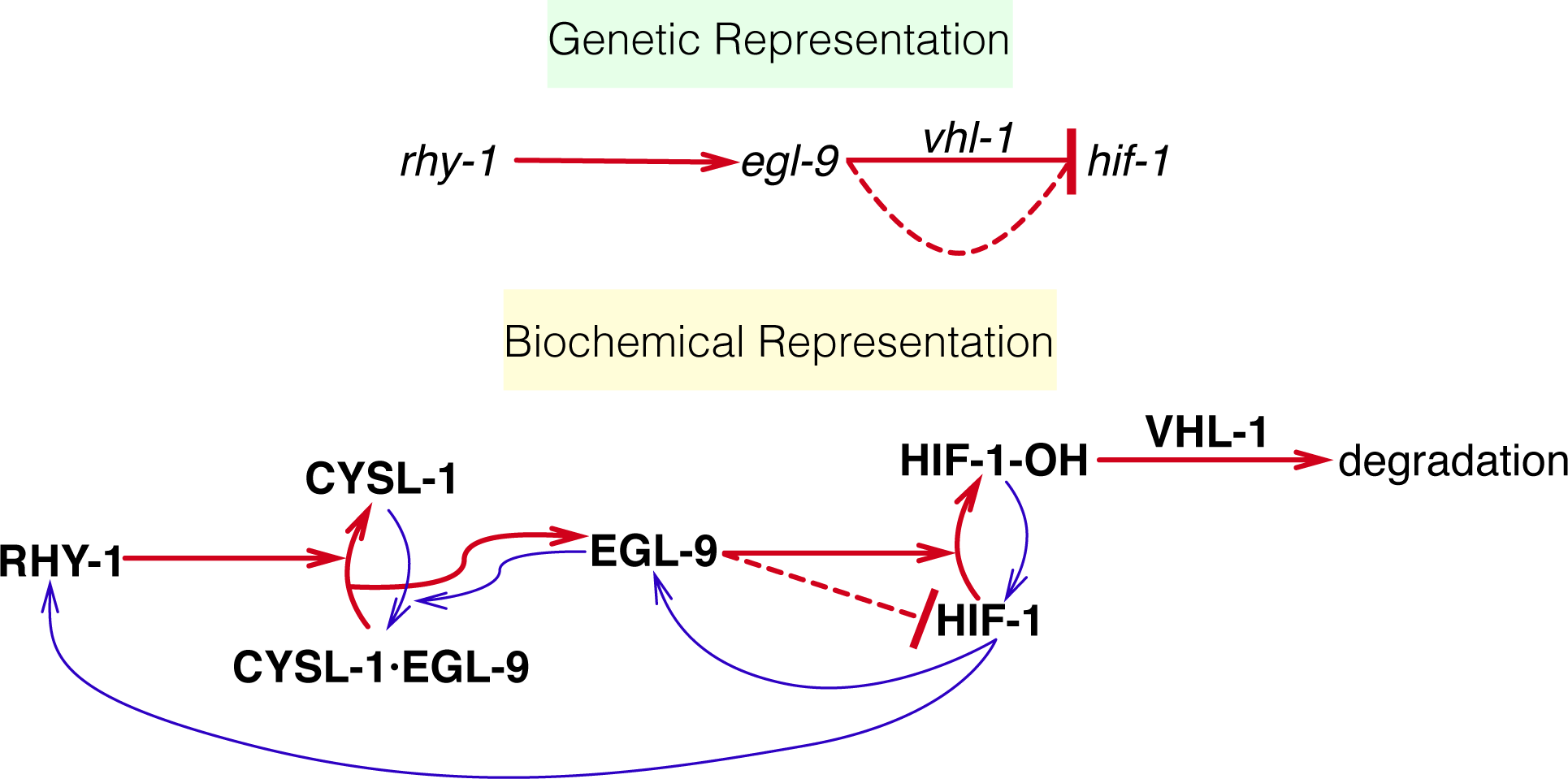
Genetic and biochemical representation of the hypoxia pathway in *C. elegans*. Red arrows are arrows that lead to inhibition of HIF-1, and blue arrows are arrows that increase HIF-1 activity or are the result of HIF-1 activity. EGL-9 is known to exert VHL-1-dependent and independent repression on HIF-1 as shown in the genetic diagram. The VHL-1-independent repression of HIF-1 by EGL-9 is denoted by a dashed line and is not dependent on the hydroxylating activity of EGL-9. Technically, RHY-1 inhibits CYSL-1, which in turn inhibits EGL-9, but this interaction was abbreviated in the genetic diagram for clarity.

Our reconstruction of the hypoxia pathway in *C. elegans* using RNA sequencing shows that whole-animal transcriptome profiles can be used as phenotypes for genetic analysis and that the phenomenon of epistasis, a hallmark of genetic interaction observed in double mutants, holds at the molecular systems level. We demonstrate that transcriptomes can aid in ordering genes in a pathway using only single mutants. Finally, we were able to identify genes that appear to be downstream of *egl-9* and *vhl-1*, but do not appear to be targets of *hif-1*. Using a single set of transcriptome-wide measurements, we observed most of the known transcriptional effects of *hif-1* as well as novel effects not described before in *C. elegans*. Taken together, this analysis demonstrates that whole-animal RNA-seq is an extremely fast and powerful method for genetic analyses in an area where phenotypic measurements are now the rate-limiting step.

## Results

### The hypoxia pathway controls thousands of genes in *C. elegans*

We selected four null single mutants within the hypoxia pathway for expression profiling: *egl-9(sa307)*, *rhy-1(ok1402)*, *vhl-1(ok161)*, *hif-1(ia4)*. We also sequenced the transcriptomes of two double mutants, *egl-9; vhl-1* and *egl-9 hif-1* as well as wild-type (N2). Each genotype was sequenced in triplicate at a depth of 15 million reads per sample. We performed whole-animal RNA-seq at a moderate sequencing depth (~7 million mapped reads per sample) under normoxic conditions. We identified around 22,000 different isoforms per sample, which allowed us to measure differential expression of 18,344 isoforms across all replicates and genotypes (~70% of the protein coding isoforms in *C. elegans*). We included in our analysis a *fog-2(q71)* mutant we have previously studied^14^, because *fog-2* is not reported to interact with the hypoxia pathway. We analyzed our data using a general linear model on logarithm-transformed counts. Changes in gene expression are reflected in the regression coefficient *β*, which is specific to each isoform within a genotype (excluding wild-type, which is used as baseline). Statistical significance is achieved when the q-value of a *β* coefficient (p-values adjusted for multiple testing) are less than 0.1. Genes that are significantly altered between the wild type and a given mutant (differentially expressed genes, DEGs) have *β* values that are statistically significantly different from 0 (i.e. greater than 0 or less than 0). *β* coefficients are analogous to the logarithm of the fold-change between the mutant and the wild type. Larger magnitudes of *β* correspond to larger perturbations (see Fig. 2). When we refer to *β* coefficients and q-values, it will always be in reference to isoforms. However, we report the sizes of each gene set in by the number of genes they contain, not isoforms. For the case of *C. elegans*, this difference is negligible since the great majority of protein-coding genes have a single isoform. We have opted for this method of referring to gene sets because it simplifies the language considerably. A complete version of the code used for this analysis with ample documentation, is available at https://wormlabcaltech.github.io/mprsq.

**Figure 2.**
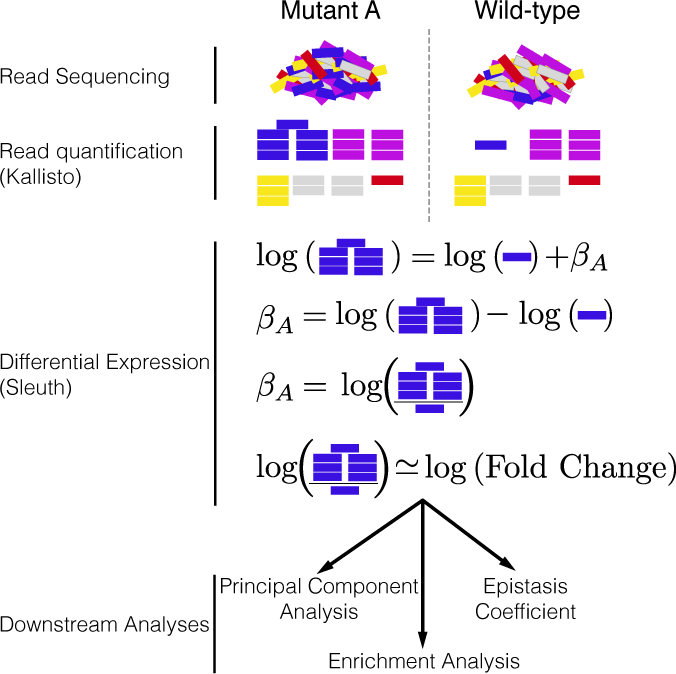
Analysis workflow. After sequencing, reads are quantified using Kallisto. Bars show estimated counts for each isoform. Differential expression is calculated using Sleuth, which outputs one *β* coefficient per isoform per genotype. *β* coefficients are analogous to the natural logarithm of the fold-change relative to a wild-type control. Downstream analyses are performed with *β* coefficients that are statistically significantly different from 0. Q-values less than 0.1 are considered statistically different from 0.

Transcriptome profiling of the hypoxia pathway revealed that this pathway controls thousands of genes in *C. elegans* (see Table 1). The *egl-9(lf)* transcriptome showed differential expression of 2,549 genes. 3,005 genes were differentially expressed in *rhy-1(lf)* mutants. The *vhl-1(lf)* transcriptome showed considerably fewer DEGs (1,275), possibly because *vhl-1* is a weaker inhibitor of *hif-1* than *egl-9*^17^. The *egl-9(lf)*;*vhl-1(lf)* double mutant transcriptome showed 3,654 DEGs. The *hif-1(lf)* mutant showed a transcriptomic phenotype involving 1,075 genes. The *egl-9(lf) hif-1(lf)* double mutant showed a similar number of genes with altered expression (744 genes).

**Table 1.** Number of differentially expressed genes in each mutant strain with respect to wild-type (N2).

| Genotype | Differentially Expressed Genes |
| --- | --- |
| *egl-9(lf)* | 2,549 |
| *rhy-1(lf)* | 3,005 |
| *vhl-1(lf)* | 1,275 |
| *hif-1(lf)* | 1,075 |
| *egl-9(lf); vhl-1(lf)* | 3,654 |
| *egl-9(lf) hif-1(lf)* | 744 |
| *fog-2(lf)* | 2,840 |

### Principal Component Analysis visualizes epistatic relationships between genotypes

PCA is used to identify relationships between high-dimensional data points^25^. We performed PCA on our data to examine whether each genotype clustered in a biologically relevant manner. PCA identifies the vector that can explain most of the variation in the data; this is called the first PCA dimension. Using PCA, one can identify the first *n* dimensions that can explain more than 95% of the variation in the data. Sample clustering in these *n* dimensions often indicates biological relationships between the data, although interpreting PCA dimensions can be difficult.

The first dimension of the PCA analysis was able to discriminate between mutants that have constitutive high levels of HIF-1 and mutants that have no HIF-1, whereas the second dimension was able to discriminate between mutants within the hypoxia pathway and outside the hypoxia pathway (see Fig. 3; *fog-2* is not reported to act in the hypoxia pathway and acts as a negative control). Therefore, expression profiling measures enough signal to cluster genes in a meaningful manner in complex metazoans.

**Figure 3.**
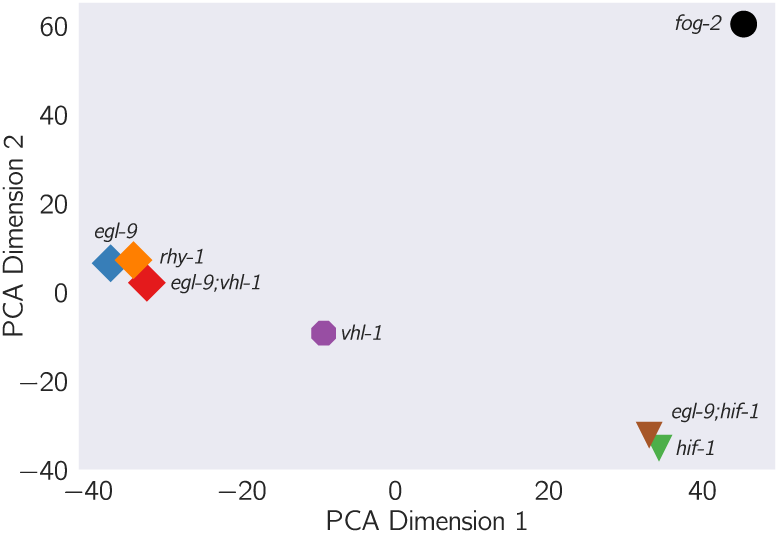
Principal component analysis of various *C. elegans* mutants. Genotypes that have an constitutive hypoxia response (i.e. *egl-9(lf)*) cluster far from genotypes that do not have a hypoxic response (i.e. *hif-1(lf)*) along PCA Dimension 1. PCA Dimension 2 separates genotypes that do not participate hypoxic response pathway.

### Reconstruction of the hypoxia pathway from first genetic principles

To reconstruct a genetic pathway, we must first assess whether two genes act on the same phenotype. If they do not act on the same phenotype (two mutations do not cause the same genes to become differentially expressed relative to wild-type), these mutants are independent. Otherwise, we must measure whether these genes act additively or epistatically on the phenotype of interest; if there is epistasis we must measure whether it is positive or negative, in order to assess whether the epistatic relationship is a genetic suppression or a synthetic interaction. To allow coherent comparisons of different mutant transcriptomes (the phenotype we are studying here), we define the shared transcriptomic phenotype between two mutants (STP) as the shared set of genes or isoforms whose expression in both mutants are different from that in wild-type, regardless of the direction of change.

### Genes in the hypoxia mutant act on the same transcriptional phenotype

All the hypoxia mutants had a significant STP: the fraction of differentially expressed genes that was shared between mutants ranged from a minimum of 10% shared between *hif-1(lf)* and *egl-9(lf); vhl-1(lf)* to a maximum of 32% shared genes between *egl-9(lf)* and *egl-9(lf); vhl-1(lf)*. For comparison, we also analyzed a previously published *fog-2(lf)* transcriptome^14^. The *fog-2* gene is involved in masculinization of the *C. elegans* germline, which enables sperm formation, and is not known to be involved in the hypoxia pathway. The hypoxia pathway mutants and the *fog-2(lf)* mutant also had STPs (8.8%–14% genes).

Next, we performed pairwise correlations between all mutant pairs. We rank-transformed the *β* coefficients of each isoform between the STP of two mutants, and calculated lines of best fit using Bayesian regressions that are robust to outliers (see Fig 4). For mutants associated with the hypoxia pathway, these correlations have values higher than 0.9 with a tight distribution around the line of best fit. The correlations for mutants from the hypoxia pathway with the *fog-2(lf)* mutant were considerably weaker, with magnitudes between 0.6–0.85 and greater variance around the line of best fit. Although *hif-1* is known to be genetically repressed by *egl-9*, *rhy-1* and *vhl-1* ^15,16^, all the correlations between mutants of these genes and *hif-1(lf)* were positive.

**Figure 4.**
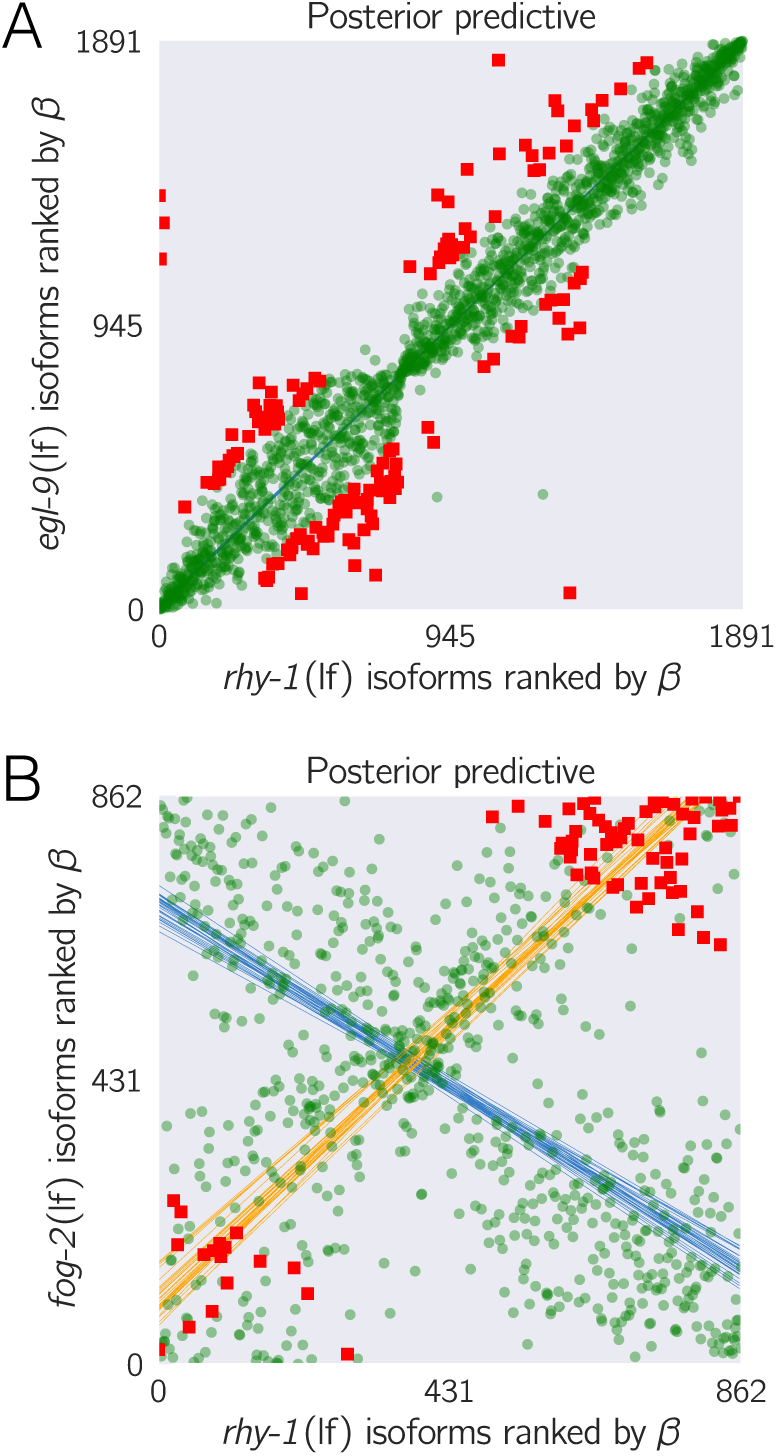
Strong transcriptional correlations can be identified between genes that share a positive regulatory connection. **A**. We obtained identified isoforms that were differentially expressed in both *egl-9(lf)* and the *rhy-1(lf)* mutants, and ranked each isoform according to its *β* coefficient. We plotted the rank of each gene in *rhy-1(lf)* versus the rank of the same gene in the *egl-9(lf)* transcriptome. **B**. For comparison, we followed the same procedure with the *fog-2(lf)* and *rhy-1(lf)* transcriptomes. *fog-2(lf)* is not known to interact with the primary hypoxia pathway. Green, transparent large points mark inliers to the primary regressions (blue lines); red squares mark outliers to the primary regressions and orange lines represent the secondary correlations involving the outliers. isoform’s x-coordinate is the sum of the regression coefficients from the single mutants *a*^−^ and *b*^−^. The Y-axis represents the deviations from the log-additive (null) model, and can be calculated as the difference between the predicted and the observed *β* coefficients. Only genes that are differentially expressed in all three genotypes are plotted. This attempts to ensure that the isoforms to be examined are regulated by both genes. These plots will generate specific patterns that can be described through linear regressions. The slope of these lines, *s*(*a*, *b*), is the transcriptome-wide epistasis coefficient.

### Transcriptome-wide epistasis

Ideally, any measurement of transcriptome-wide epistasis should conform to certain expectations. First, it should make use of the regression coefficients of as many genes as possible. Second, it should be summarizable in a single, well-defined number. Third, it should have an intuitive behavior, such that special values of the statistic have an unambiguous interpretation.

We found an approach that satisfies all of the above conditions and which can be graphed in a plot we call an epistasis plot (see Fig 5) In an epistasis plot, the X-axis represents the expected *β* coefficient for given gene in a double mutant *a*^−^*b*^−^ if *a* and *b* interact log-additively. In other words, each individual isoform’s x-coordinate is the sum of the regression coefficients from the single mutants *a*^−^ and *b*^−^. The Y-axis represents the deviations from the log-additive (null) model, and can be calculated as the difference between the predicted and the observed *β* coefficients. Only genes that are differentially expressed in all three genotypes are plotted. This attempts to ensure that the isoforms to be examined are regulated by both genes. These plots will generate specific patterns that can be described through linear regressions. The slope of these lines, *s*(*a*, *b*), is the transcriptome-wide epistasis coefficient.

**Figure 5.**
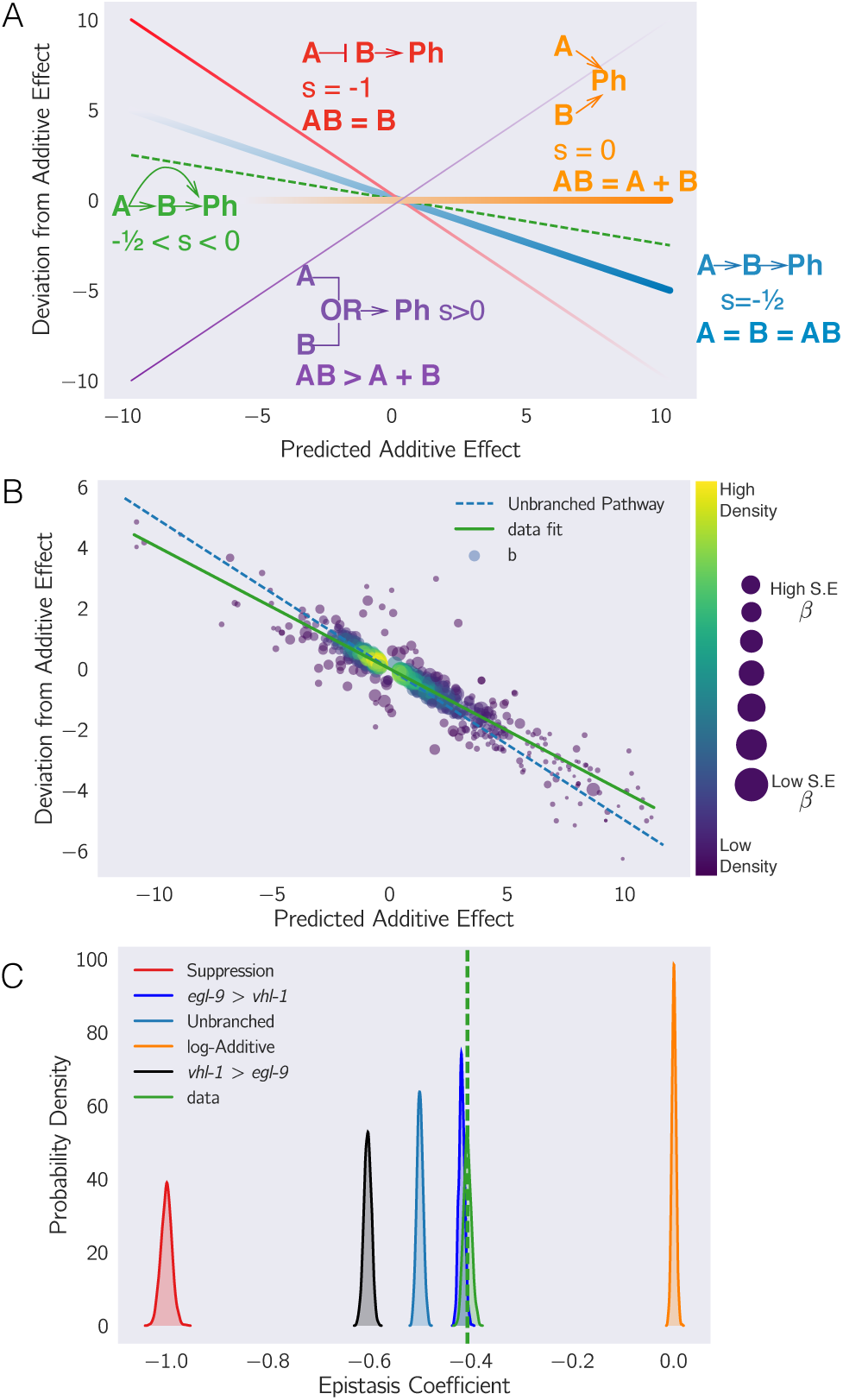
(**A**) Schematic diagram of an epistasis plot. The X-axis on an epistasis plot is the expected coefficient for a double mutant under an log-additive model (null model). The Y-axis plots deviations from this model. Double mutants that deviate in a systematic manner from the null model exhibit transcriptome-wide epistasis (*s*). To measure *s*, we find the line of best fit and determine its slope. Genes that act log-additively on a phenotype **(Ph)** will have *s* = 0 (null hypothesis, orange line); whereas genes that act along an unbranched pathway will have *s* = −1/2 (blue line). Strong repression is reflected by *s* = −1 (red line), whereas *s* > 0 correspond to synthetic interactions (purple line). (**B**) Epistasis plot showing that the *egl-9(lf); vhl-1(lf)* transcriptome deviates significantly from a null additive. Points are colored qualitatively according to density (purple—low, yellow—high) and size is inversely proportional to the standard error (S.E.) of the y-axis. The green line is the line of best fit from an orthogonal distance regression. (**C**) Comparison of simulated epistatic coefficients against the observed coefficient. Green curve shows the bootstrapped observed transcriptome-wide epistasis coefficient for *egl-9* and *vhl-1*. Dashed green line shows the mean value of the data. Simulations use only the single mutant data to idealize what expression of the double mutant should look like. *a* > *b* means that the phenotype of *a* is observed in a double mutant *a*^−^*b*^−^.

Transcriptome-wide epistasis coefficients can be understood intuitively for simple cases of genetic interactions if true genetic nulls are used. If two genes act additively on the same set of differentially expressed isoforms then all the plotted points will fall along the line *y* = 0. If two genes act positively in an unbranched pathway, then all the mutants should have the same phenotype. It follows that data from this pathway will form line with slope equal to 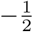. On the other hand, in the limit of complete genetic inhibition of *b* by *a* in an unbranched pathway (i.e., *a* is in great excess over *b*, such that under the conditions measured *b* has no activity), the plots should show a line of best fit with slope equal to −1. Genes that interact synthetically (*i.e.*, through an OR-gate) will fall along lines with slopes > 0. When there is epistasis of one gene over another, the points will fall along one of two possible slopes that must be determined empirically from the single mutants data. We can use both single mutants data to predict the distribution of slopes that results for the cases stated above. The transcriptome-wide epistasis coefficient emerges as a powerful way to quantify epistasis because it integrates information from many different isoforms into a single number (see Fig. 5).

In our experiment, we studied two double mutants, *egl-9(lf) hif-1(lf)* and *egl-9(lf); vhl-1(lf)*. We wanted to understand how well an epistatic analysis based on transcriptome-wide coefficients agreed with the epistasis results reported in the literature, which were based on qPCR of single genes. Therefore, we performed orthogonal distance regression on the two gene combinations we studied (*egl-9* and *vhl-1*, and *egl-9* and *hif-1*) to determine the epistasis coefficient for each gene pair. We also generated models for the special cases mentioned above using the single mutant data.

We measured the epistasis coefficient between *egl-9* and *vhl-1*: *s*(*egl-9 vhl-1*) = −0.41. Simulations using just the single mutant data showed that the double mutant exhibited the *egl-9(lf)* phenotype (see Fig. 5). We used Bayesian model selection to reject a linear pathway (odds ratio (OR) > 10^83^), which leads us to conclude *egl-9* is upstream of *vhl-1* acting on a phenotype in a branched manner. We also measured epistasis between *egl-9* and *hif-1*, *s*(*egl-9*, *hif-1*) = −0.80, and we found that this behavior could be predicted by modeling *hif-1* downstream of *egl-9*. We also rejected the null hypothesis that these two genes act in a positive linear pathway (OR> 10^93^). Taken together, this leads us to conclude that *egl-9* strongly inhibits *hif-1*.

### Epistasis can be predicted

Given our success in measuring epistasis coefficients, we wanted to know whether we could predict the epistasis coefficient between *egl-9* and *vhl-1* in the absence of the *egl-9(lf)* genotype. Since RHY-1 indirectly activates EGL-9, the *rhy-1(lf)* transcriptome should contain more or less equivalent information to the *egl-9(lf)* transcriptome. Therefore, we generated predictions of the epistasis coefficient between *egl-9* and *vhl-1* by substituting in the *rhy-1(lf)* data, predicting *s*(*rhy* − 1, *vhl* − 1) = −0.45. Similarly, we used the *egl-9(lf); vhl-1(lf)* double mutant to measure the epistasis coefficient while replacing the *egl-9(lf)* dataset with the *rhy-1(lf)* dataset. We found that the epistasis coefficient using this substitution was −0.40. This coefficient was different from −0.50 (OR > 10^62^), reflecting the same qualitative conclusion that *vhl-1* represents a branch in the hypoxia pathway. In conclusion, we were able to obtain a quantitatively close prediction of the epistasis coefficient for two mutants using the transcriptome of a related, upstream mutant. Finally, in the absence of a single mutant, an upstream locus can be used to estimate epistasis between two genes.

### Transcriptomic decorrelation can be used to infer functional distance

So far, we have shown that RNA-seq can accurately measure genetic interactions. However, genetic interactions do not require two gene products to interact biochemically, nor even to be physically close to each other. RNA-seq cannot measure physical interactions between genes, but we wondered whether expression profiling contains sufficient information to order genes along a pathway.

Single genes are often regulated by multiple independent sources. The connection between two nodes can in theory be characterized by the strength of the edges connecting them (the thickness of the edge); the sources that regulate both nodes (the fraction of inputs common to both nodes); and the genes that are regulated by both nodes (the fraction of outputs that are common to both nodes). In other words, we expected that expression profiles associated with a pathway would respond quantitatively to quantitative changes in activity of the pathway. Targeting a pathway at multiple points would lead to expression profile divergence as we compare nodes that are separated by more degrees of freedom, reflecting the flux in information between them.

We investigated this possibility by weighting the robust Bayesian regression between each pair of genotypes by the size of the shared transcriptomic phenotype of each pair divided by the total number of isoforms differentially expressed in either mutant (*N*_Intersection_/*N*_Union_). We plotted the weighted correlation of each gene pair, ordered by increasing functional distance (see Fig. 6). In every case, we see that the weighted correlation decreases monotonically due mainly, but not exclusively, to a smaller STP.

**Figure 6.**
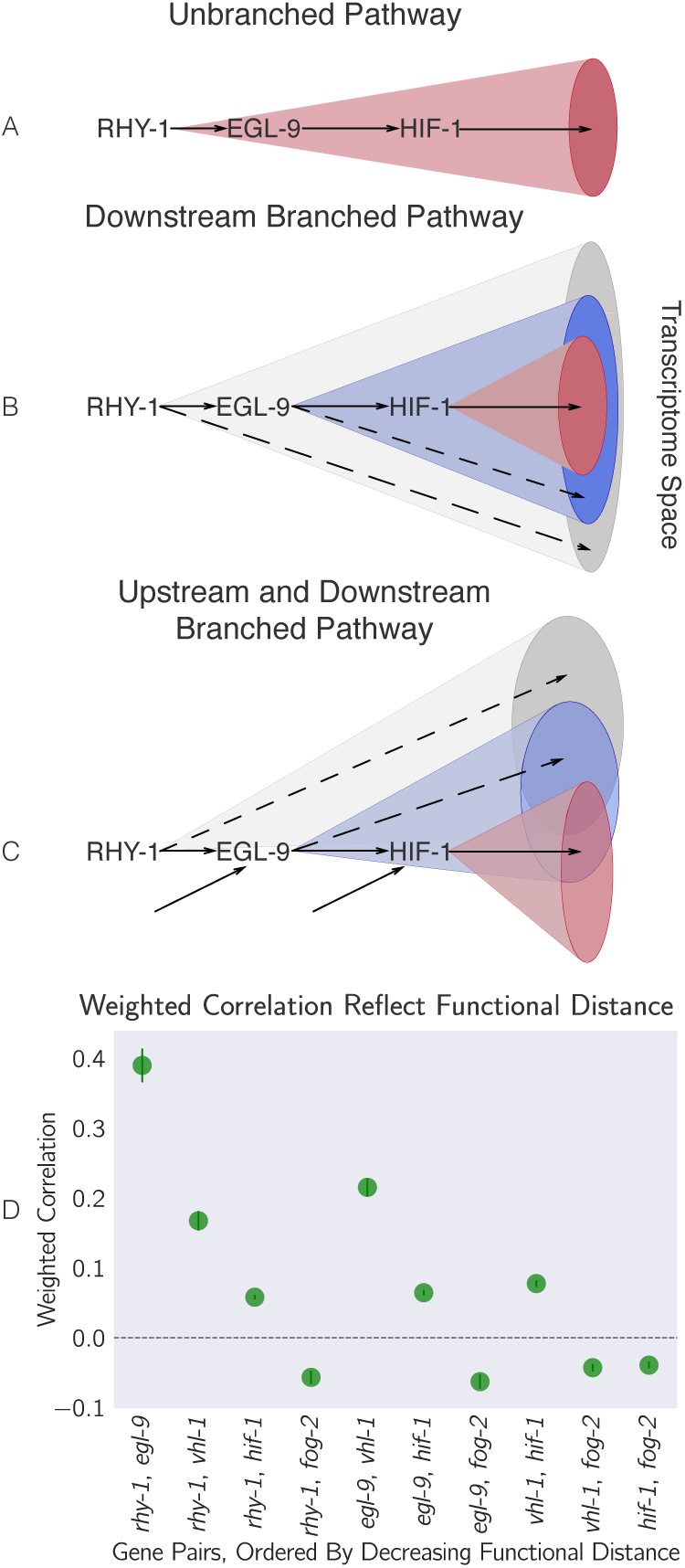
Theoretically, transcriptomes can be used to order genes in a pathway under certain assumptions. Arrows in the diagrams above are intended to show the direction of flow, and do not indicate valence. **A**. A linear pathway in which *rhy-1* is the only gene controlling *egl-9*, which in turn controls *hif-1* does not contain information to infer the order between genes. **B**. If *rhy-1* and *egl-9* have transcriptomic effects that are separable from *hif-1*, then the *rhy-1* transcriptome should contain contributions from *egl-9*, *hif-1* and *egl-9* - and *hif-1*-independent pathways. This pathway contains enough information to infer order. **C**. If a pathway is branched both upstream and downstream, transcriptomes will show even faster decorrelation. Nodes that are separated by many edges may begin to behave almost independently of each other with marginal transcriptomic overlap or correlation. **D**. The hypoxia pathway can be ordered. We hypothesize the rapid decay in correlation is due to a mixture of upstream and downstream branching that happens along this pathway. Bars show the standard error of the weighted coefficient from the Monte Carlo Markov Chain computations.

We believe that this result is not due to random noise or insufficiently deep sequencing. Instead, we propose a framework in which every gene is regulated by multiple different molecular species, which induces progressive decorrelation. This decorrelation in turn has two consequences. First, decorrelation within a pathway implies that two nodes may be almost independent of each other if the functional distance between them is large. Second, it may be possible to use decorrelation dynamics to infer gene order in a branching pathway, as we have done with the hypoxia pathway.

### 0.1 Classical epistasis identifies a core hypoxic response

We identified a main hypoxia response induced by HIF-1 (466 genes) by selecting genes that were consistently altered in *egl-9(lf)*, *rhy-1(lf)*, *vhl-1(lf)* and *egl-9(lf); vhl-1(lf)* mutants but which were suppressed in the *egl-9(lf) hif-1(lf)* mutant. This response included five transcription factors (*W02D7.6*, *nhr-57*, *ztf-18*, *nhr-135* and *dmd-9*; Supplementary Table 1). Even though HIF-1 is an activator, not all of these genes were up-regulated. We reasoned that only genes that are up-regulated in HIF-1-inhibitor mutants are candidates for direct regulation by HIF-1. We found 264 such genes. Phenotype Enrichment Analysis^26^ showed that this gene list was enriched in genes associated with oxygen response, dauer development and dauer constitutive phenotypes (fold-change > 4 and *q* < 10^−1^ for all terms).

### Feedback can be inferred

While some of the rank plots contained a clear positive correlation (see Fig. 4), others showed a discernible cross-pattern (see Figure S2). In particular, this cross-pattern emerged between *vhl-1(lf)* and *rhy-1(lf)* or between *vhl-1(lf)* and *egl-9(lf)*, even though *vhl-1*, *rhy-1* and *egl-9* are all inhibitors of *hif-1(lf)*. Such cross-patterns could be indicative of feedback loops or other complex interaction patterns. If the above is correct, then it should be possible to identify genes that are regulated by *rhy-1* in a logically consistent way: Since loss of *egl-9* causes *rhy-1* mRNA levels to increase, if this increase leads to a significant change in RHY-1 activity, then it follows that the *egl-9(lf)* and *rhy-1(lf)* should show anti-correlation in a subset of genes. Since we do not observe many genes that are anti-correlated, we conclude that is unlikely that the change in *rhy-1* mRNA expression causes a significant change in RHY-1 activity under normoxic conditions. We also searched for genes with *hif-1*-independent, *vhl-1*-dependent gene expression and found 71 genes (Supplementary Table 2).

### Identification of non-classical epistatic interactions

*hif-1(lf)* has traditionally been viewed as existing in a genetic OFF state under normoxic conditions. However, our dataset indicates that 1,075 genes show altered expression when *hif-1* function is removed in normoxic conditions. Moreover, we observed positive correlations between *hif-1(lf) β* coefficients and *egl-9(lf)*, *vhl-1(lf)* and *rhy-1(lf) β* coefficients in spite of the negative regulatory relationships between these genes and *hif-1*. Such positive correlations could indicate a relationship between these genes that has not been reported previously.

We first identified genes that exhibited violations of the canonical genetic model of the hypoxia pathway. We searched for genes that changed in different directions between *egl-9(lf)* and *vhl-1(lf)*, or, equivalently, between *rhy-1(lf)* and *vhl-1(lf)* (we assume that all results from the *rhy-1(lf)* transcriptome reflect a complete loss of *egl-9* activity) without specifying any further conditions. We found 56 that satisfied this condition (see Fig. 7, Supplementary Table 3). When we checked expression of these genes in the double mutant, we found that *egl-9* remained epistatic over *vhl-1* for this class of genes. This class of genes may in fact be much larger because it overlooks genes that have wild-type expression in an *egl-9(lf)* background, altered expression in a *vhl-1(lf)* background, and suppressed (wild-type) expression in an *egl-9(lf); vhl-1(lf)* background.

**Figure 7.**
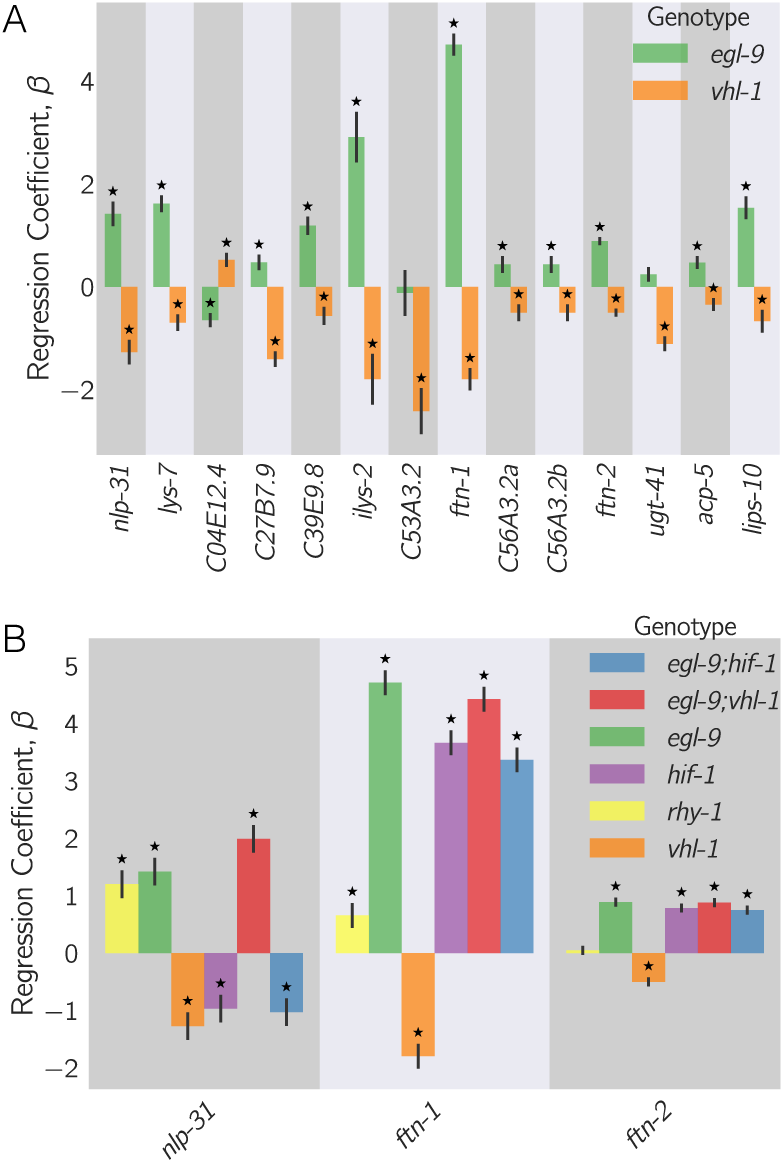
**A.** 56 genes in *C. elegans* exhibit non-classical epistasis in the hypoxiapathway, characterized by opposite effects on gene expression, relative to the wild-type, of of the *vhl-1(lf)* compared to *egl-9(lf)* (or *rhy-1(lf)*) mutants. Shown are a random selection of 15 out of 56 genes for illustrative purposes. **B**. Genes that behave non-canonically have a consistent pattern. *vhl-1(lf)* mutants have an opposite effect to *egl-9(lf)*, but *egl-9* remains epistatic to *vhl-1* and loss-of-function mutations in *hif-1* suppress the *egl-9(lf)* phenotype. Asterisks show *β* values significantly different from 0 relative to wild-type (*q* < 10^−1^).

Although this entire class had similar behavior, for simplicity we focused on two genes, *nlp-31* and *ftn-1* which have expression patterns representative of this class. *ftn-1* is described to be responsive to mutations in the hypoxia pathway and has been reported to have aberrant behaviors; specifically, mutation of *egl-9* and *vhl-1* have opposing effects on *ftn-1* expression^27,28^. These studies showed the same patterns of *ftn-1* expression phenotypes using both RNAi and alleles, which allays concerns of strain-specific interference. We observed that *hif-1* was epistatic to *egl-9*, and that *egl-9* and *hif-1* both promoted *ftn-1* expression.

Analysis of *ftn-1* expression reveals that *egl-9* is epistatic to *hif-1*; that *vhl-1* has opposite effects to *egl-9*, and that *vhl-1* is epistatic to *egl-9*. Analysis of *nlp-31* reveals similar relationships. *nlp-31* expression is decreased in *hif-1(lf)*, and increased in *egl-9(lf)*. However, *egl-9* is epistatic to *hif-1*. Like *ftn-1*, *vhl-1* has the opposite effect to *egl-9*, yet is epistatic to *egl-9*. We propose in the Discussion a novel model for how HIF-1 might regulate these targets.

## Discussion

### The *C. elegans* hypoxia pathway can be reconstructed *de novo* from RNA-seq data

In this paper, we have shown that whole-organism transcriptomic phenotypes can be used to reconstruct genetic pathways and to discern previously overlooked or uncharacterized genetic interactions. We successfully reconstructed the hypoxia pathway, and inferred order of action (*rhy-1* activates *egl-9*, *egl-9* and *vhl-1* inhibit *hif-1*), and we were able to infer from transcriptome-wide epistasis measurements that *egl-9* exerts *vhl-1*-dependent and independent inhibition on *hif-1*.

### Interpretation of the non-classical epistasis in the hypoxia pathway

The observation of 56 genes that exhibit a specific pattern of non-classical epistasis suggests the existence of previously undescribed aspects of the hypoxia pathway. Some of these non-classical behaviors had been observed previously^27,28,29^, but no satisfactory mechanism has been proposed to explain this biology. Previous studies^28,27^ suggested that HIF-1 integrates information on iron concentration in the cell to determine its binding affinity to the *ftn-1* promoter, but could not definitively establish a mechanism. It is unclear why deletion of *hif-1* and deletion of *egl-9* both cause induction of *ftn-1* expression, but deletion of *vhl-1* abolishes this induction. Moreover, Luchachack et al^29^ have previously reported that certain genes important for the *C. elegans* immune response against pathogens reflect similar non-canonical expression patterns. Their interpretation was that *swan-1*, which encodes a binding partner to EGL-9^30^, is important for modulating HIF-1 activity in some manner. The lack of a conclusive double mutant analysis in this work means the role of SWAN-1 in modulation of HIF-1 activity remains to be demonstrated. Other mechanisms, such as tissue-specific differences in the pathway^31^ could also modulate expression, though it is worth pointing out that *ftn-1* expression appears restricted to a single tissue, the intestine^32^. Another possibility is that *egl-9* controls *hif-1* mRNA stability via other *vhl-1*-independent pathways, but we did not see a decreases in *hif-1* level in *egl-9(lf)*, *rhy-1(lf)* or *vhl-1(lf)* mutants. Another possibility, such as control of protein stability via *egl-9* independently of *vhl-1* ^33^ will not lead to splitting unless it happens in a tissue-specific manner.

One parsimonious solution is to consider HIF-1 as a protein with both activating and inhibiting states. In fact, HIF-1 already exists in two states in *C. elegans*: unmodified HIF-1 and HIF-1-hydroxyl (HIF-1-OH). Under this model, the effects of HIF-1 for certain genes like *ftn-1* or *nlp-31* are antagonized by HIF-1-hydroxyl, which is present at only a low level in the cell in normoxia because it is degraded in a *vhl-1*-dependent fashion. This means that loss of *vhl-1* stabilizes HIF-1-hydroxyl. Genes that are sensitive to HIF-1-hydroxyl will be inhibited as a result of the increase in the amount of this HIF-1-hydroxyl, despite loss of *vhl-1* function also increasing the level of non-hydroxylated HIF-1. On the other hand, *egl-9(lf)* abrogates the generation of HIF-1-hydroxyl, stimulating accumulation of non-hydroxylated HIF-1 and promoting gene expression. Whether deletion of *hif-1(lf)* is overall activating or inhibiting will depend on the relative activity of each protein state under normoxia (see Fig. 8). HIF-1-hydroxyl is challenging to study genetically, and if it does have the activity suggested by our genetic evidence this may have prevented such a role from being detected. No known mimetic mutations are available with which to study the pure hydroxylated HIF-1 species, and mutations in the Von Hippel-Lindau gene that stabilize the hydroxyl species also increase the quantity of non-hydroxylated HIF-1 by mass action.

**Figure 8.**
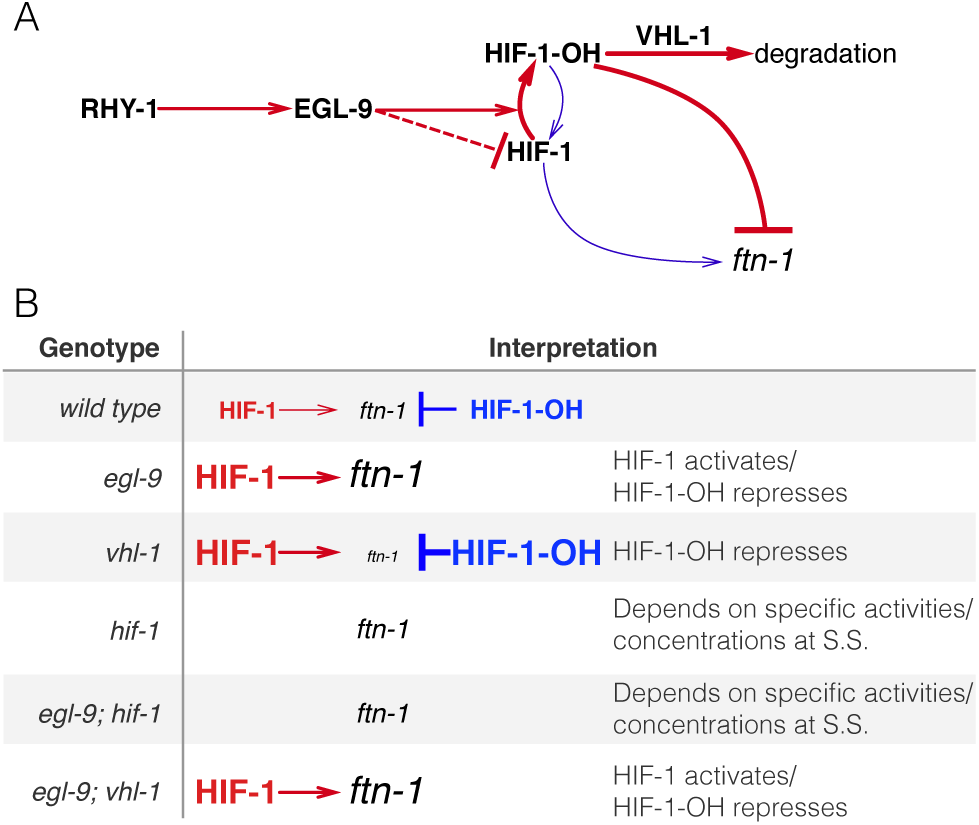
A hypothetical model showing a mechanism where HIF-1-hydroxyl antagonizes HIF-1. **A**. Diagram showing that RHY-1 activates EGL-9. EGL-9 hydroxylates HIF-1 in an oxygen-dependent fashion. Under normoxia, HIF-1 is rapidly hydroxylated and only slowly does hydroxylated HIF-1 return to its original state. EGL-9 can also inhibit HIF-1 in an oxygen-independent fashion. HIF-1-OH is rapidly degraded in a VHL-1-dependent fashion. In our model, HIF-1 and HIF-1-OH have opposing effects on transcription. The width of the arrows represents the rates under normoxic conditions. **B**. Table showing the effects of loss-of-function mutations on HIF-1 and HIF-1-OH activity, showing how this can potentially explain the *ftn-1* expression levels in each case. S.S = Steady-state.

Because HIF-1 is detected at low levels in cells under normoxic conditions^34^, total HIF-1 protein levels are assumed to be so low as to be biologically inactive. However, our data show 1,075 genes change expression in response to loss of *hif-1* under normoxic conditions, which establishes that there is sufficient total HIF-1 protein to be biologically active. Our analyses also revealed that *hif-1(lf)* shares positive correlations with *egl-9(lf)*, *rhy-1(lf)* and *vhl-1(lf)*, and that each of these genotypes also shows a secondary negative rank-ordered expression correlation with each other.

A homeostatic argument can be made in favor of the activity of HIF-1-hydroxyl. The cell must continuously monitor multiple metabolite levels. The *hif-1*-dependent hypoxia response integrates information from O_2_, *α*-ketoglutarate and iron concentrations in the cell. One way to integrate this information is by encoding it within the effective hydroxylation rate of HIF-1 by EGL-9. Then the dynamics in this system will evolve exclusively as a result of the total amount of HIF-1 in the cell. Such a system can be sensitive to fluctuations in the absolute concentration of HIF-1^35^. Since the absolute levels of HIF-1 are low in normoxic conditions, small fluctuations in protein copy-number can represent a large fold-change in HIF-1 levels. These fluctuations would not be problematic for genes that must be turned on only under conditions of severe hypoxia—presumably, these genes would be associated with low affinity sites for HIF-1, so they are only activated when HIF-1 levels are far above random fluctuations, such as in hypoxia.

For yet other sets of genes that must change expression in response to the hypoxia pathway, it may not be sufficient to integrate metabolite information exclusively via EGL-9-dependent hydroxylation of HIF-1. In particular, genes that may function to increase survival in mild hypoxia may benefit from regulatory mechanisms that can sense minor changes in environmental conditions and which therefore benefit from robustness to transient changes in protein copy number. Likewise, genes that are involved in iron or *α*-ketoglutarate metabolism (such as *ftn-1*) may benefit from being able to sense, accurately, small and consistent deviations from basal concentrations of these metabolites. For these genes, the information may be better encoded by using HIF-1 and HIF-1-hydroxyl as an activator/repressor pair. Such circuits are known to possess distinct advantages for controlling output in a manner that is robust to transient fluctuations in the levels of their components^36,37^.

Our RNA-seq data suggests that one of these atypical targets of HIF-1 may be RHY-1. Although *rhy-1* does not exhibit non-classical epistasis, all genotypes containing a *hif-1(lf)* mutation had increased expression levels of *rhy-1*. We speculate that if *rhy-1* is controlled by both HIF-1 and HIF-1-hydroxyl, then this might imply that HIF-1 regulates the expression of its pathway (and therefore itself) in a manner that is robust to total HIF-1 levels.

### Insights into genetic interactions from vectorial phenotypes

Here, we have described a set of methods that can be applied to any vectorial phenotype studied with an appropriate experimental design. Transcriptome profiling methods offers a lot of information, but transcriptome-wide interpretation of the results is often extremely challenging. Each method has its own advantages and disadvantages.

Principal component analysis is computationally tractable and clusters can often be visually detected with ease. However, PCA can be misleading, especially when the dimensions represented do not explain a very large fraction of the variance present in the data. In addition, principal dimensions are the product of a linear combination of vectors, and therefore must be interpreted with extreme care. In this case, the first principal dimension separated genotypes that increase HIF-1 protein levels from those that decrease it. Although PCA showed that there is information hidden in these genotypes, it is not enough by itself to provide biological insight.

Whereas PCA operates on all genotypes simultaneously, correlation analysis is a pairwise procedure that measures how predictable the gene expression changes are in a mutant given the vector of expression changes in another. Like PCA, correlation analysis is easy and fast. Unlike PCA, the product of a correlation analysis is a single number with a straightforward interpretation. However, correlation analysis is sensitive to outliers. Although outliers can be mitigated via rank-transformations, these transformations cannot remove outliers resulting from systematic variation caused, for example, by feedback loops. Such interactions can lead to vanishing correlations if both are equally strong. Adequately weighted correlations could be informative for ordering genes along pathways. A drawback of correlation analysis is that the number of pairwise comparisons increases combinatorially.

Another way to analyze genetic interactions is via general linear models (GLMs) that include interaction terms between two or more genes. GLMs can quantify the genetic interactions on single genes. We and others^13,14^ have used GLMs to perform epistasis analyses of a pathway using transcriptomic phenotypes. GLMs are powerful, but they generate a different interaction coefficient for each gene measured. The large number of coefficients makes interpretation of the genetic interaction between two mutants extremely difficult. Previous approaches^13^ have attempted to visualize these coefficients via heatmaps.

The epistasis plots we demonstrate here are a novel way to visualize epistasis in vectorial phenotypes. Here, we have shown how an epistasis plot can be used to identify interactions between two genes by examining the transcriptional phenotypes of single mutants and the double mutant. In reality, epistasis plots can be generated for any set of measurements involving a set of *N* mutants (or conditions) and an *N*-mutant genotype. Epistasis plots can accumulate an arbitrary number of points within them, possess a rich structure that can be visualized and have straightforward interpretations for special slope values. Epistasis plots and GLMs are not antagonistic toward one another. Indeed, one could use a GLM to quantify interactions at single-gene resolution, then plot the conglomerated results in an epistasis plot instead of as a heatmap (for a non-genetic example, see^14^).

Until relatively recently, the rapid generation and molecular characterization of null mutants was a major bottleneck for genetic analyses. Advances in genomic engineering mean that, for a number of organisms, production of mutants is now rapid and efficient. As mutants become easier to produce, biologists are realizing that phenotyping and characterizing the biological functions of individual genes is challenging. This is particularly true for whole organisms, where subtle phenotypes can go undetected for long periods of time. We have shown that whole-animal RNA-sequencing is a sensitive method that can be seamlessly incorporated with genetic analyses of epistasis.

## Materials and Methods

### Strains

Strains used were N2 wild-type (Bristol), JT307 *egl-9(sa307)*, CB5602 *vhl-1(ok161)*, ZG31 *hif-1(ia4)*, RB1297 *rhy-1(ok1402)*. CB6088 *egl-9(sa307) hif-1(ia4)* CB6116 *egl-9(sa307)*;*vhl-1(ok161)*, All lines were grown on standard nematode growth media (NGM) Petri plates seeded with OP50 *E. coli* at 20°C^38^.

### RNA isolation

Lines were synchronized by harvesting eggs via sodium hypochlorite treatment and subsequently plating eggs on food. Worms were staged and based on the time after plating, vulva morphology and the absence of eggs. 30–50 non-gravid young adults were picked and placed in 100 μL of TE pH 8.0 (Ambion AM9849) in 0.2 mL PCR tubes on ice. Worms were allowed to settle or spun down by centrifugation and ~ 80 μL of supernatant removed before flash-freezing in liquid *N*_2_. These samples were digested with Proteinase K (Roche Lot No. 03115 838001 Recombinant Proteinase K PCR Grade) for 15 min at 60° in the presence of 1% SDS and 1.25 μL RNA Secure (Ambion AM7005). RNA samples were then taken up in 5 Volumes of Trizol (Tri-Reagent Zymo Research) and processed and treated with DNase I using Zymo MicroPrep RNA Kit (Zymo Research Quick-RNA MicroPrep R1050). RNA was eluted in RNase-free water stored at −80°C. Samples were analyzed using a NanoDrop (Thermo Fisher) for impurities, Qubit for concentration and then analyzed on an Agilent 2100 BioAnalyzer (Agilent Technologies). Replicates were selected that had RNA integrity numbers (RIN) equal or greater than 9.0 and showed no evidence of bacterial ribosomal bands, except for the ZG31 mutant where one of three replicates had a RIN of 8.3.

### Library preparation and sequencing

10 ng of quality checked total RNA from each sample was reverse-transcribed into cDNA using the Clontech SMARTer Ultra Low Input RNA for Sequencing v3 kit (catalog #634848) in the SMARTSeq2 protocol^39^. RNA was denatured at 70°C for 3 min in the presence of dNTPs, oligo dT primer and spiked-in quantitation standards (NIST/ERCC from Ambion, catalog #4456740). After chilling to 4°C, the first-strand reaction was assembled using the LNA TSO primer described in Picelli et al^39^, and run at 42°C for 90 minutes, followed by denaturation at 70°C for 10 min. The entire first strand reaction was then used as template for 13 cycles of PCR using the Clontech v3 kit. Reactions were cleaned up with 1.8x volume of Ampure XP SPRI beads (catalog #A63880) according to the manufacturer’s protocol. After quantification using the Qubit High Sensitivity DNA assay, a 3 ng aliquot of the amplified cDNA was run on the Agilent HS DNA chip to confirm the length distribution of the amplified fragments. The median value for the average cDNA lengths from all length distributions was 1,076 bp. Tagmentation of the full length cDNA for sequencing was performed using the Illumina/Nextera DNA library prep kit (catalog #FC-121–1030). Following Qubit quantitation and Agilent BioAnalyzer profiling, the tagmented libraries were sequenced. Libraries were sequenced on Illumina HiSeq2500 in single read mode with the read length of 50 nt to an average depth of 15 million reads per sample following manufacturer’s instructions. Base calls were performed with RTA 1.13.48.0 followed by conversion to FASTQ with bcl2fastq 1.8.4. Spearman correlation of the estimated counts for each genotype showed that every pairwise correlation within genotype was 0.9.

### Read alignment and differential expression analysis

We used Kallisto^40^ to perform read pseudo-alignment and performed differential analysis using Sleuth^41^. We fit a general linear model for a transcript *t* in sample *i*:
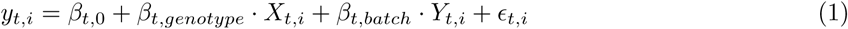

where *y_t_*_,*i*_ was the logarithm transformed counts; *β_t_*_,*genotype*_ and *β_t_*_,*batch*_ were parameters of the model, and which could be interpreted as biased estimators of the log-fold change; *X_t_*_,*i*_, *Y_t_*_,*i*_ were indicator variables describing the conditions of the sample; and *ϵ_t_*_,*i*_ was the noise associated with a particular measurement.

### Genetic Analysis, Overview

The processed data were analyzed using Python 3.5. We used the Pandas, Matplotlib, Scipy, Seaborn, Sklearn, Networkx, PyMC3, and TEA libraries^42,43,44,45,46,47,48,49^. Our analysis is available in Jupyter Notebooks50. All code and processed data are available at https://github.com/WormLabCaltech/mprsq along with version-control information. Our Jupyter Notebook and interactive graphs for this project can be found at https://wormlabcaltech.github.io/mprsq/. Raw reads were deposited in the Short Read Archive under the study accession number SRP100886.

### Weighted correlations

Pairwise correlations between transcriptomes were calculated by identifying the set of DEGs common to both transcriptomes under analysis. DEGs were rank-ordered according to their regression coefficient, *β*. Bayesian robust regressions were performed using a Student-T distribution using the PyMC3 library^45^ (pm.glm.families.StudenT in Python). If the correlation had an average value > 1, the average correlation coefficient was set to 1. Weights were calculated as the proportion of genes that were inliers to a regression divided by the total number of DEGs present in either mutant.

### Epistatic analysis

The epistasis coefficient between two null mutants *a* and *b* was calculated as:
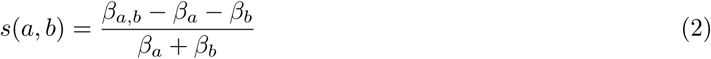

Null models for various epistatic relationships were generated by sampling the single mutants in an appropriate fashion. For example, to generate the distribution for two mutants that obey the epistatic relationship *a*^−^ = *a*^−^*b*^−^, we substituted *β_a_*_,*b*_ with *β_a_* and bootstrapped the result.

To select between theoretical models, we implemented an approximate Bayesian Odds Ratio. We defined a free-fit model, *M*_1_, that found the line of best fit for the data:
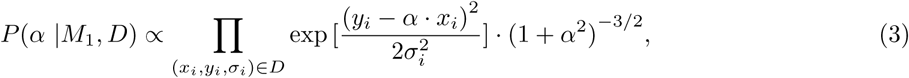

where *α* was the slope to be determined, *x_i_*, *y_i_* are the of each point, and *σ_i_* was the standard error associated with the y-value. We used equation 3 to obtain the most likely slope given the data, *D*, via minimization (scipy.optimize.minimize in Python). Finally, we approximated the odds ratio as:
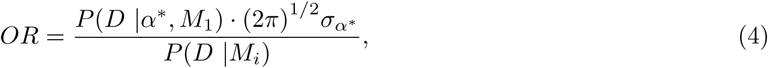

where *α*^*^ was the slope found after minimization, 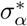 was the standard deviation of the parameter at the point *α*^*^ and *P*(*D* |*M_i_*) was the probability of the data given the parameter-free model, *M_i_*.

### Enrichment analysis

Tissue, Phenotype and Gene Ontology Enrichment Analysis were carried out using the WormBase Enrichment Suite for Python^26,48^.

## Acknowledgements

This work was supported by HHMI with whom PWS is an investigator and by the Millard and Muriel Jacobs Genetics and Genomics Laboratory at California Institute of Technology. All strains were provided by the CGC, which is funded by NIH Office of Research Infrastructure Programs (P40 OD010440). This article would not be possible without help from Dr. Igor Antoshechkin who performed all sequencing. We thank Hillel Schwartz, Jonathan Liu, Han Wang, and Porfirio Quintero and Erich Schwarz for their advice, support and conversations throughout this project.

